# Inflammation cellular platform (INCEPLAT) for testing anti-inflammatory compounds for SARS-CoV-2

**DOI:** 10.1101/2024.09.25.614910

**Authors:** Blanca D. López-Ayllón, Laura Mendoza-García, Ana de Lucas-Rius, Tránsito García-García, Raúl Fernández-Rodríguez, Antonio Romero-Guillén, Judit Serrat, Sonia Zúñiga, Natalia Redondo, Mar Siles-Lucas, Juan J. Garrido, María Montoya

## Abstract

From the early days of the COVID-19 pandemic, an excessive release of proinflammatory cytokines, such as IL6, was detected in serum from patients. As a consequence, several anti-inflammatory drugs, such as Dexamethasone (a strong corticoid), were used to counteract such cytokine storm occurring during severe disease. By contrast, pro-inflammatory interleukin 11 (IL11), a member of the IL6 family, was detected in respiratory tissues from infected patients and in experimental epithelial cellular models. In this work, human A549 lung epithelial cells were individually transduced with SARS-CoV-2 open reading frames (ORFs), resulting in a IL11 increase, which was significantly decreased after Dexamethasone treatment. The use of this cellular platform allowed us to screen for new possible anti-inflammatory compounds from *Fasciola hepatica*. Our results highlighted the ability of FhNEJ (*Fasciola hepatica* newly excysted juvenile flukes) somatic extract to decrease IL11 levels in ORF-transduced cells. These results emphasized the role of IL11 in lung epithelial inflammation, making it a potential target for future treatments of lung inflammation which occurs in COVID-19, and validate the use of these ORF-expressing cells as a cellular platform to test anti-inflammatory compounds for COVID-19 disease.

## INTRODUCTION

COVID-19 (COronaVIrus Disease-2019) is a potentially fatal respiratory disease caused by the Severe Acute Respiratory Syndrome Coronavirus 2 (SARS-CoV-2), which rapidly spread worldwide in 2020. Since the very beginning of the pandemic, it was evident that aggressive and uncontrolled inflammatory responses were part of the problem in some infected patients, resulting in damage to the airways. Like the rest of Coronaviruses, SARS-CoV-2 genome consists of a single-stranded positive-sense RNA molecule of approximately 29,900 nucleotides (NCBI Reference Sequence: NC_045512.2) arranged into 13 open reading frames (ORFs) and encoding 27 proteins ^1^. Among them, there are seven ORFs that encode accessory proteins: ORF3a, ORF3c, ORF6, ORF7a, ORF7b, ORF8 and ORF9b, and additional ORFs which have been suggested but it remains controversial (ORF3b, ORF3d, ORF9c and ORF10) ^2–6^. As their name suggest, accessory proteins are dispensable for viral replication, but several reports have demonstrated their involvement in COVID-19 pathogenesis by mediating antiviral host responses ^7–17^. SARS-CoV-2 mostly affects the respiratory tract usually leading to pneumonia in most patients, and to Acute Respiratory Distress Syndrome (ARDS) in 15% of cases. ARDS is mainly triggered by elevated levels of pro-inflammatory cytokines in serum, such as Interleukin 6 (IL6) ^18,19^, referred to as cytokine storm ^20^. Interleukin 11 (IL11), a member of the IL6 family of cytokines, is similar to IL6 and both form a GP130 heterodimer complex to initiate its downstream signaling ^21–24^, but their respective hexameric signaling complex formation differ ^25^. In addition, while IL6R (interleukin 6 receptor) is expressed most highly on immune cells, IL11RA (interleukin 11 receptor subunit alpha) is expressed in stromal cells, such as fibroblasts and hepatic stellate cells, and also on parenchymal cells, including hepatocytes. Hence, it may be expected that IL6 biology relates mostly to immune functions whereas IL11 activity is more closely linked to the stromal and parenchymal biology ^21,25^. Since the nineties, high IL11 release during viral infections have been described ^26–28^, and more recently, several studies have related this interleukin to fibrosis, chronic inflammation and matrix extracellular remodeling^10,21,25,29–32^. Among the anti-inflammatory drugs used since the emergence of the pandemic, Dexamethasone (a strong corticoid), was one of the most commonly used to counteract the cytokine storm occurring during severe COVID-19 ^33,34^.

*Fasciola hepatica* is a helminth parasite with an indirect life cycle involving water snails as intermediary hosts and mammals, typically ruminants and humans, as definitive hosts ^35^. *F. hepatica*, and generally all parasitic helminths, have evolved sophisticated mechanisms that allow them to escape from host immunity ^36,37^. Such mechanisms induce an overall state of immune down-regulation and tolerance that has been suggested to mitigate the immune pathology that is associated to hyper-inflammatory conditions such as allergies, autoimmune diseases and, most recently, SARS-CoV-2 infection ^38–43^.

In this study, A549 lung epithelial cells were individually transduced with open reading frames: ORF3a, ORF3d, ORF6, ORF7a, ORF7b, ORF8, ORF9b, ORF9c or ORF10 from SARS-CoV-2 (Wuhan-Hu-1 isolate). Then, pro-inflammatory interleukins were detected, and IL11 was found mainly upregulated, which was dramatically reduced by Dexamethasone treatment. We named this cellular platform INCEPLAT (Inflammation Cellular PLATform). Thus, INCEPLAT led us to evaluate the potential anti-inflammatory activity of tegument- and somatic-enriched protein extracts of newly excysted juveniles of *Fasciola hepatica* (FhNEJ-Teg and FhNEJ-Som, respectively), as a proof-of-concept of our cellular platform, showing that this system was suitable for investigating compounds with potential anti-inflammatory activity against SARS-CoV-2-induced IL-11 secretion (Figure 1).

**Figure 1.**
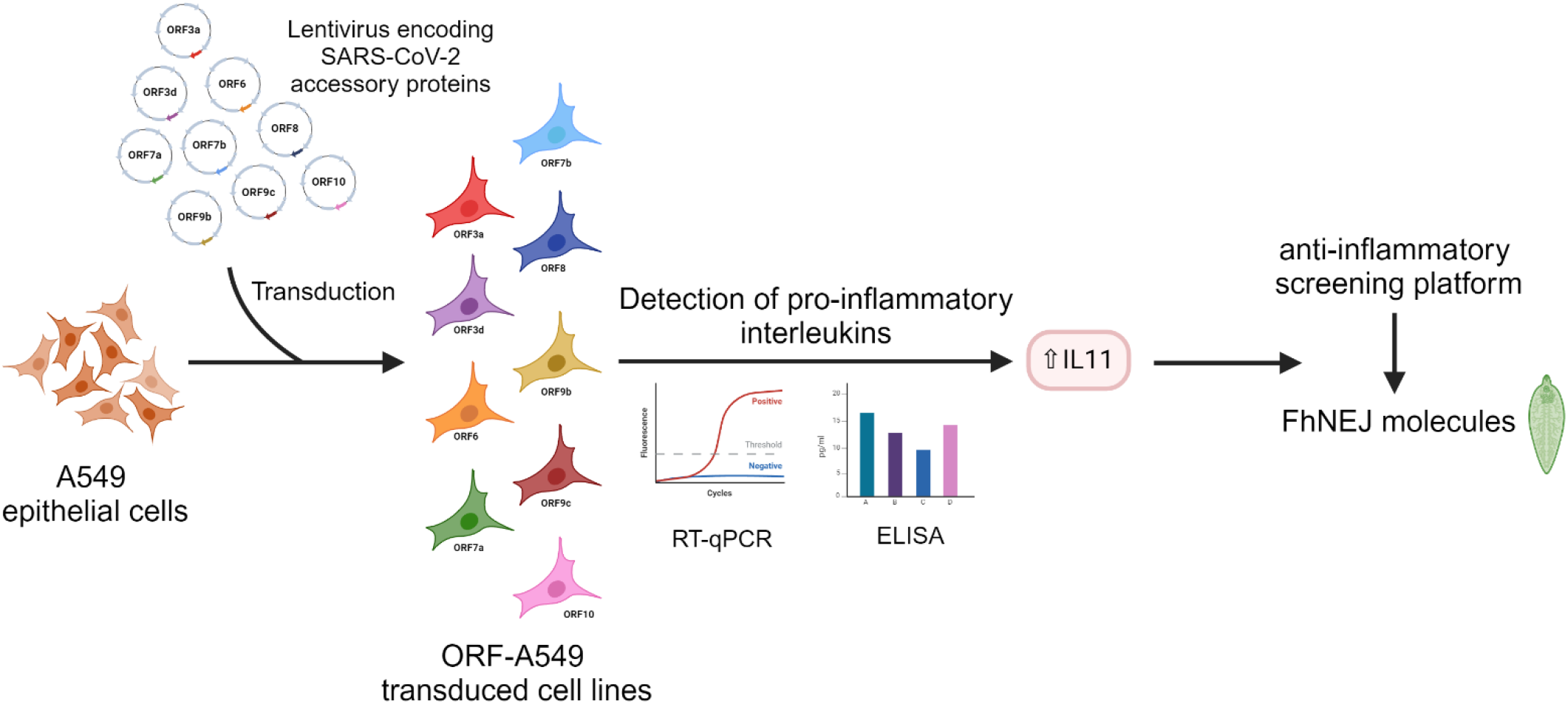
Experimental workflow. Created with BioRender.com. FhNEJ: *Fasciola hepatica* newly excysted juvenile flukes.

## MATERIAL & METHODS

### Cell culture, lentivirus production and transduction

A549 pulmonary epithelial cells (ATCC CRM-CCL-185; RRID: CVCL_0023) were cultured in Dulbecco’s Modified Eagle Medium (DMEM) (Gibco, #41966029) supplemented with 10% (v/v) heat-inactivated fetal bovine serum (FBS) (Gibco, #1027016), 1% Penicillin-Streptomycin (100U/ml) (Gibco, #15070063) and Amphotericin B (Gibco, #15290026). ORF3a, ORF3d, ORF6, ORF7a, ORF7b, ORF8, ORF9b, ORF9c or ORF10 accessory proteins coding sequences (codon-optimized for mammalian expression) were cloned into pLVX-EF1α-IRES-Puro Cloning and Expression Lentivector (Clontech, Takara, #631253) to generate pseudotyped lentiviral particles encoding each accessory protein of SARS-CoV-2 (Wuhan-Hu-1 isolate) at the CNIC (Centro Nacional de Investigaciones Cardiovasculares) Viral Vector Unit (ViVU) as described previously ^9–11^. Accessory proteins were C-terminally 2xStrep-tagged to check viral protein expression. A549 cells were transduced by incubating them with lentivirus at a MOI of 10 for 24 h followed by 2 µg/ml puromycin treatment to start the selection of successfully transduced cells. All cells were cultured at 37°C and 5% CO_2_.

### RNA isolation

Cells were seeded (3 x 10^5^) in 6-well plates and lysed using RLT buffer for RNA isolation (RNeasy Mini kit, Qiagen, #74106). Each sample was performed in triplicate. RNA was isolated following the manufactureŕs protocol, quantified by nanodrop 1000 (Thermo Scientific) and quality controlled by Bioanalyzer (Agilent).

### Real time qPCR

RNA samples (500 ng) were reverse transcribed using qScript™ cDNA synthesis kit (Quanta Biosciences Inc., #95047), following manufacturer’s instructions. Primers sequences are available in Supplementary Table 1 (Table S1). The final 15 µL PCR reaction included 2 μL of 1:5 diluted cDNA as template, 3 µL of 5x PyroTaq EvaGreen qPCR Mix Plus with ROX (Cultek Molecular Bioline, #88H24), and transcript-specific forward and reverse primers at a 20 μM final concentration. Real time PCR was carried out in a QuantStudio 12K Flex system (Applied Biosystems) under the following conditions: 15 min at 95 °C followed by 40 cycles of 30 s at 94 °C, 30 s at 57 °C and 45 s at 72 °C. Melting curve analyses were performed at the end, to ensure specificity of each PCR product. Relative expression results were calculated using GenEx6 Pro software (MultiD-Göteborg, Sweden), based on the Cq values obtained.

### SARS-CoV-2 infection

Confluent A549-ACE2 cells (kindly provided by R. Andino, University of California San Francisco, USA), seeded in 24-well plates were mock infected or infected with SARS-CoV-2 isolate ^15^ at a MOI of 1. Total RNA was extracted at 24 hpi using RNeasy Mini kit (Qiagen, #74106) following manufacturer’s specifications. To decrease sample variability, nine independent infections were performed in each case; and for subsequent RT-qPCR analysis, RNAs were pooled three by three, obtaining three biological replicates for each condition ^44^.

### ELISA experiments and Dexamethasone treatment

Cells were seeded (6 x 10^4^) in 24-well plates with 1 ml of medium, and cell supernatants were collected after 24 h. Interleukins levels were detected with Human IL-6 Uncoated ELISA Kit (Invitrogen, #88-7066-88), Human IL-8 Uncoated ELISA Kit (Invitrogen, #88-8086-88), and Human IL-11 DuoSet ELISA Kits (RD Systems, #DY218) according to the manufacturer’s instructions. To perform Dexamethasone (Sigma, #D4902) treatment, cells were cultured as indicated above, and treated with 10 µM of Dexamethasone. After 24 h supernatants were collected and IL11 levels were detected by ELISA.

### Fasciola hepatica protein extracts

Tegument- and somatic-enriched antigenic extracts of *F. hepatica* newly excysted juveniles (FhNEJ-Teg and FhNEJ-Som, respectively) were obtained by stimulating the excystment of *F. hepatica* metacercariae (Ridgeway Research) *in vitro*, as previously described^45^. Briefly, metacercariae were first incubated for one hour at 37 °C in a CO_2_-supplemented solution containing 0.02 M sodium dithionite (Sigma, #15795-3), washed twice in distilled water and resuspended in 5 ml of Hank’s balanced salt solution (Sigma, # H9394) supplemented with 10% lamb bile (obtained from a local abattoir) and 30 mM HEPES (Sigma, #H3375) at pH 7.4. Metacercariae were incubated for up to five hours at 37 °C and FhNEJ were manually recovered every hour after addition of excystment media and left to recover for at least one hour at 37°C and 5% CO_2_ in RPMI-1640 culture media (Thermo Fisher Scientific, #31870-017) containing 30 mM HEPES (Sigma, #H3375), 0.1% glucose (Sigma, #G7021) and 50 µg/ml gentamycin (Sigma, #G1914). After recovery, FhNEJ were washed twice in PBS and FhNEJ-Teg proteins were extracted with an appropriate volume of PBS containing 1% of Nonidet-P40 (Sigma, #74385). Following incubation for thirty minutes at room temperature (RT) and mild rotation, FhNEJ were centrifuged for five minutes at 300 xg and the supernatant containing the FhNEJ-Teg protein fraction was transferred to a clean tube. FhNEJ-Som was obtained by resuspending pellets containing naked FhNEJ in an appropriate volume of RIPA buffer (Sigma, #R0278) followed by incubation on ice for fifteen minutes, sonication (five cycles of thirty seconds on/thirty seconds off) and clarification by centrifugation for ten minutes at 13000 xg and 4°C. To avoid cell toxicity, detergents in FhNEJ-Teg and FhNEJ-Som were removed as previously described ^46^: FhNEJ-Teg was incubated with SM-2 Adsorbent Bio-Beads (BioRad, #1523920) for two hours at RT with mild rotation and proteins in FhNEJ-Som were acetone-precipitated using the ReadyPrep 2-D Cleanup kit (BioRad, #1632130) according to the manufacturer’s instructions. After precipitation, FhNEJ-Som protein pellets were resuspended in 50 mM ammonium bicarbonate (Sigma, #207861) and 1% sodium deoxycholate (Acros Organics, #218590250) ^47^ to ensure solubilization of hydrophobic proteins. Protein extracts were obtained in the absence of protease and phosphatase inhibitors and stored at -80 °C until use. Protein concentrations were determined using the Pierce BCA Protein Assay kit (Thermo Fisher Scientific, #23227).

### Quantification and Statistical Analysis

Statistical analyses were performed using GraphPad Prism 5 Software. P-values were determined using Student’s t-test. Unless otherwise stated, data are shown as the mean of at least three biological replicates. Significant differences are indicated as: * = p <0.05; ** = <0.01; *** = p<0.001, **** = p<0.0001.

In Figures 4B and 4C, data plotting and statistical analyses were performed with GraphPad Prism 10 Software. Unpaired t-tests were used to compare two groups, and comparisons between more than two groups was performed using a one-way Analysis of Variance (ANOVA) test followed by Tukey post-hoc analysis for pair-wise comparisons.

## RESULTS

### Expression of SARS-CoV-2 accessory proteins increase pro-inflammatory interleukins gene expression

It is known that SARS-CoV-2 virus uses different strategies to interplay and alter the host cellular machinery. To explore the function of individual SARS-CoV-2 accessory proteins in such interaction, A549 human lung carcinoma cells were lentivirus transduced expressing individual open reading frames (ORFs) encoding such accessory proteins. These were ORF3a, ORF3d, ORF6, ORF9a, ORF7b, ORF8, ORF9b, ORF9c or ORF10 (named ORF-A549 thereafter). They were expressed with a C-terminally 2xStrep-tag to facilitate detection after expression. GFP-lentivirus-transduced or non-transduced A549 cells were used as controls in each experiment (A549 control cells), both giving the same results. ORF overexpression in A549 transduced cells was verified by immunofluorescence staining using anti-StrepTag antibody, and was previously published for ORF3a, ORF6, ORF7a, ORF7b, ORF8, ORF9b, ORF9c and ORF10 ^9,11,48^. ORF3d expression is represented in Figure S1. We observed different patterns of localization in A549 cells, as well as variable levels of ORF expression between cells in each cell line, being all positive as compared with the control counterparts.

RT-qPCR was used to assess pro-inflammatory interleukins 6 (IL6), 11 (IL11), 8 (IL8) and 1beta (IL1b) gene expression in ORF-A549 cell lines. Although IL6 appears mostly elevated in the serum of patients with COVID-19^18^, we did not detect a significant increase in IL6 expression in any of the ORF-A549 cell lines (data not shown). However, we found significantly upregulated IL11, a member of the IL6 family of cytokines, in cells expressing ORF3d, ORF7a, ORF7b, ORF9b and ORF9c (Figure 2A).

**Figure 2.**
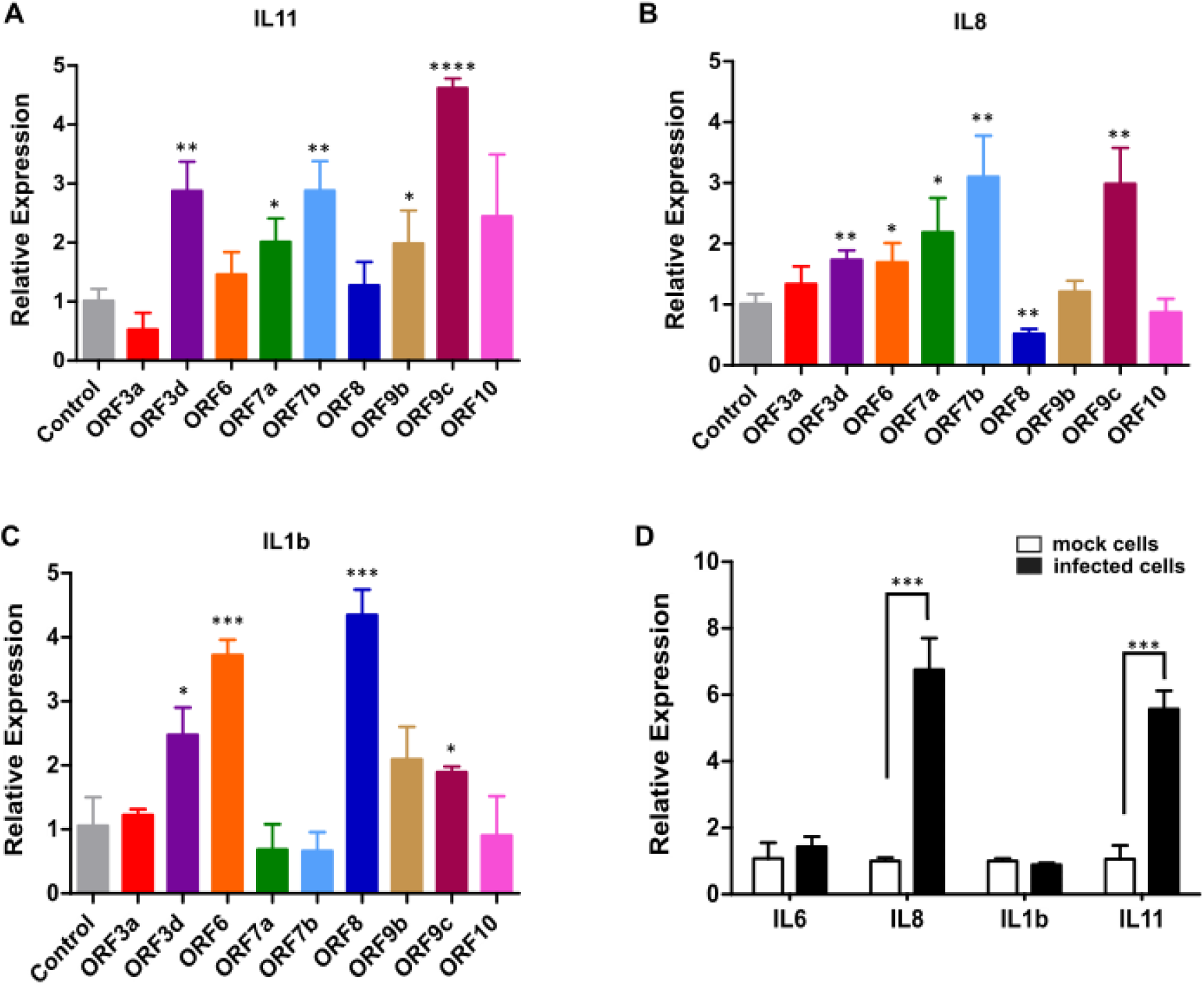
Relative expression of interleukins IL11 mRNA **(A),** IL8 **(B)** and IL1β **(C)** in A549 cells expressing SARS-CoV-2 accessory proteins by RT-qPCR. **(D)** Relative expression of IL11, IL8, IL1β and IL6 RNA in A549-ACE2 cells infected with SARS-CoV-2 virus compared to mock infected cells. Error bars represent mean ± SD (n ≥ 3). Statistical significance is given as follows: *p < 0.05, **p < 0.01, ***p < 0.001, **** p < 0.0001 compared to A549 control/mock cells.

IL8 was also significantly overexpressed in ORF3d, ORF6, ORF7a, ORF7b and ORF9c, but significantly reduced in ORF8-expressing cells (Figure 2B). Likewise, IL1b was significant overexpressed in cells expressing ORF3d, ORF6, ORF8 and ORF9c (Figure 2C). To evaluate the expression of these interleukins in the infection context, A549-hACE2 cells were infected with SARS-CoV-2 virus, and RT-qPCR for these interleukins was also performed. In line with our previous findings, increment on IL6 gene expression in infected cells was not observed (Figure 2D). Surprisingly, in contrast to IL1b expression found in ORF-A549 cell lines, IL1b overexpression was not induced by SARS-CoV-2 infection. However, IL8 and IL11 expression showed a significant increase in infected cells (Figure 2D).

### Expression of SARS-CoV-2 accessory proteins increase pro-inflammatory interleukins release

After detecting an increase in some pro-inflammatory cytokines’ gene expression, interleukins release was measured by ELISA in ORF-A549 cells. Surprisingly, we found a slightly increase of IL6 in cells expressing ORF3d, ORF8, ORF9b and ORF9c (Figure 3C). However, levels higher of 100 pg/ml of IL11 and IL8 were found in most of the transduced cell lines (Figures 3A and 3B) as compared to IL6, where we only detected levels under 60 pg/ml (Figure 3C). Dexamethasone is a potent steroid drug that affects the immune system, and it has been widely used in conditions with inflammation such as cancer, acute asthma, arthritis, Crohn’s disease or COVID-19^49–52^. Thus, we treated our ORF-A549 with Dexamethasone, and IL11 release was measured. After 24h treatment, a significant decrease in IL11 levels was observed in all treated cells compared to untreated cells (Figure 3D). By contrast, we did not observe that effect in control cells. These results show that our ORF-A549 revealed an increase in IL11 that could be reduced using anti-inflammatory drugs, paving the way for testing new compounds with potential anti-inflammatory properties in the context of SARS-CoV-2 infection.

**Figure 3.**
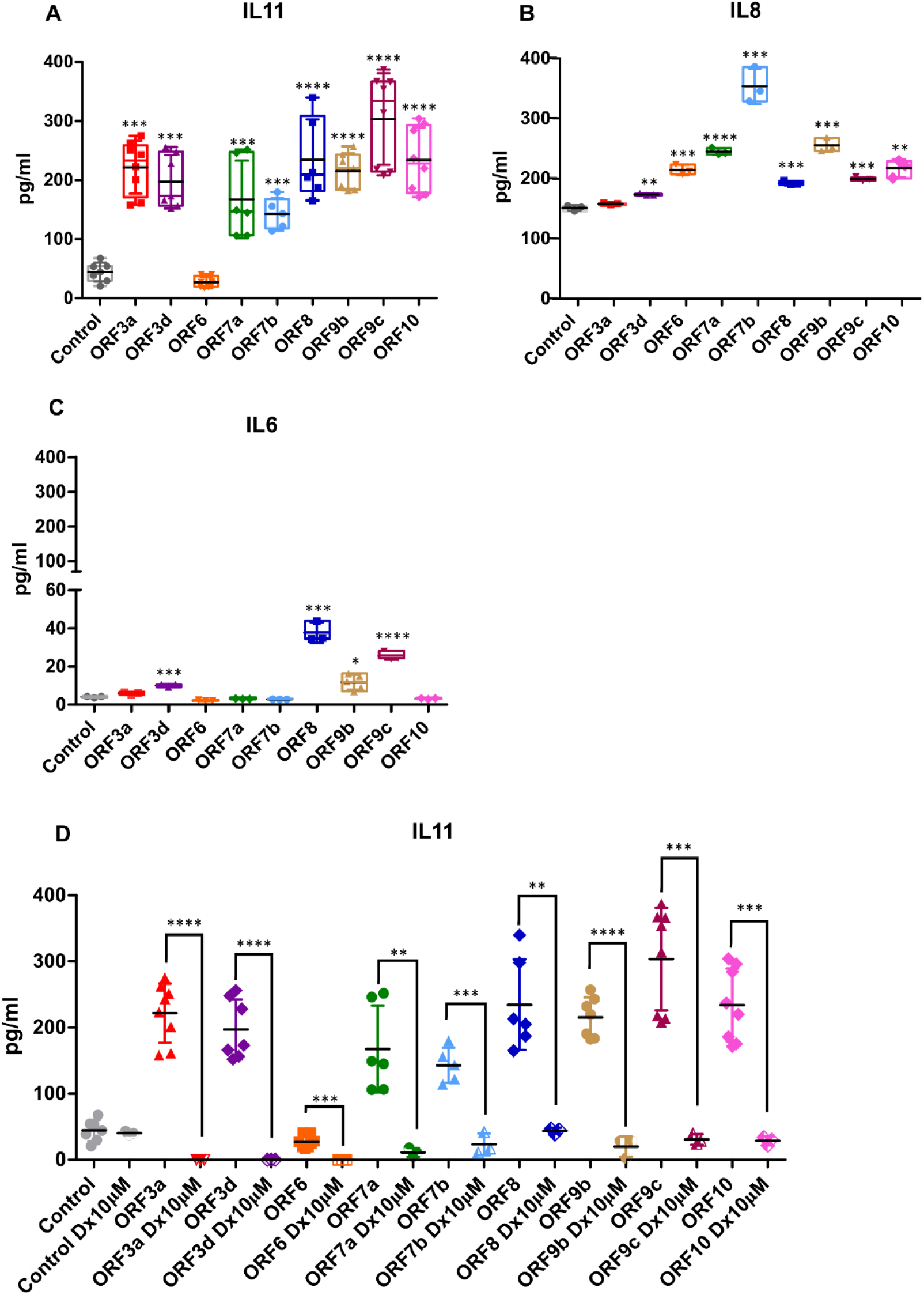
Protein measured by ELISA of secreted IL11 **(A)**, IL8 **(B)** and IL6 **(C)** by A549 transduced cells after 24h. **(D)** IL11 release after 24h treatment with 10 µM of Dexamethasone (Dx10µM). Statistical significance is given as follows: *p < 0.05, **p < 0.01, ***p < 0.001 and ****p < 0.0001 to A549 control cells, or to untreated cells in (D).

### IL11 release in ORF10-transduced A549 cells serve as a suitable platform to study the anti-inflammatory potential of novel compounds

ORF10-transduced cells were treated with tegument- and somatic-enriched protein extracts of FhNEJ (Figure 4A), which preferentially contain proteins located at the parasite surface or internal organs, respectively. IL11 secretion and cell viability were analyzed 24 hours after stimulation (Figure 4B and 4C). As expected, treatment with the positive control, Dexamethasone, resulted in the highest reduction of IL11 secretion by ORF10-transduced cells. Our results also showed that treatment with FhNEJ-Som elicited a discrete but significant inhibition of IL11 release compared to untreated ORF10-transduced control cells or those treated with the FhNEJ-Som buffer (Figure 4B), despite a decrease in cell viability in FhNEJ-Som- and Som-buffer-treated cells due to toxicity derived from buffer compounds (Figure 4C). These results sharply contrasted with those obtained from cells treated with FhNEJ-Teg, which caused no change in IL11 secretion at any of the tested concentrations.

**Figure 4.**
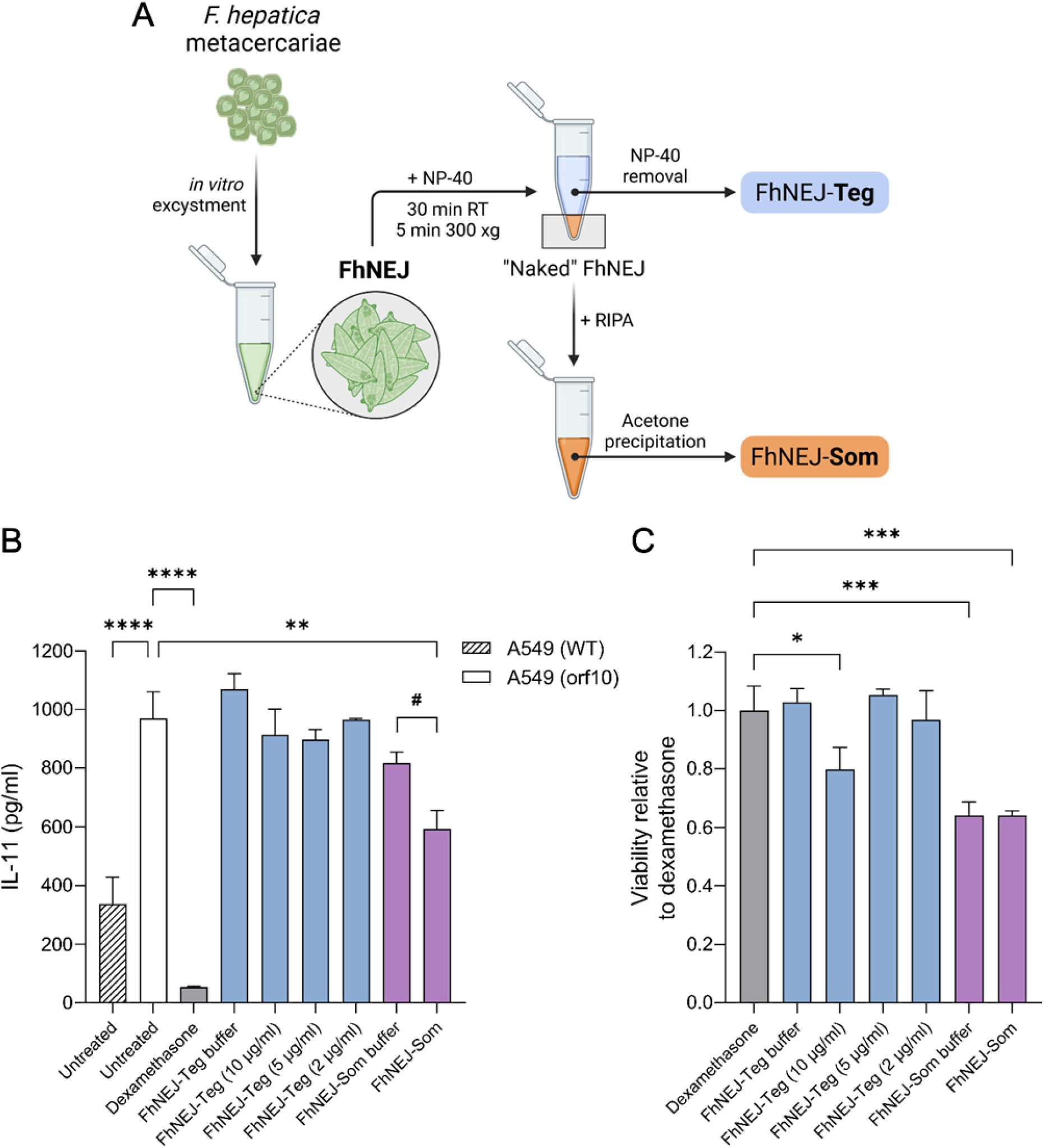
Treatment of ORF10-transduced cells with FhNEJ-Som decreases IL11 secretion. **(A)** Schematic overview of the methodology employed to obtain tegument- and somatic-enriched antigenic extracts of FhNEJ (FhNEJ-Teg and FhNEJ-Som, respectively). Created with BioRender.com. FhNEJ: *Fasciola hepatica* newly excysted juvenile flukes. A549 cells were treated with FhNEJ-Som (3.4 µg/ml) or FhNEJ-Teg at different concentrations, and IL11 secretion **(B)** and cell viability **(C)** were assessed 24 hours after treatment. Untreated cells were incubated with regular cell medium and cells treated with dexamethasone (10 µM) were used as positive controls for the inhibition of IL11 secretion. Cells treated with FhNEJ-Teg/Som buffers were included to control that the observed effects were not caused by buffer compounds. Bars indicate the mean ± SEM of three biological replicates calculated relative to ORF10-transduced cells untreated **(B)** or to dexamethasone-treated cells **(C)**. Asterisks indicate significant differences between groups (**p<0.01, ****p<0.0001, one-way ANOVA; #p<0.05, Student’s t-test). Notably, there were no significant differences in IL11 secretion between untreated orf10-transduced cells and those treated with FhNEJ-Som buffer (B), nor in cell viability between ORF10-transduced cells treated with FhNEJ-Som and those treated with FhNEJ-Som buffer (C) (Student’s t-test).

## DISCUSSION

The results of this work highlight the role of IL11 in lung epithelial inflammation, making it a potential target for future treatments of lung inflammation which takes place in COVID-19. Our results validate the use of these ORF-expressing cells as a cellular platform to test anti-inflammatory compounds for COVID-19 disease. ARDS in severe cases of COVID-19 is mainly triggered by elevated levels of pro-inflammatory cytokines, such as IL6 ^20^. IL11 is a pro-inflammatory interleukin, a member of the IL6 family of cytokines, which have been involved in numerous inflammatory processes, including viral infections ^10,11,20,25,28,29,53,54^. Indeed, it was originally discovered as a molecule in fibroblast supernatants that stimulated proliferation, but it has been also described that human alveolar and bronchial epithelial-like cells produce IL-11 ^54^. These findings correlate with our results, since IL11 was overexpressed in most A549-transduced epithelial cells, but we did not observe overexpression of IL6 in ORF-A549 cell lines (data not shown). On the other hand, it is known the involvement of IL1b and IL8 in a variety of inflammatory diseases ^55–57^. We found IL8 significantly overexpressed in ORF3d, ORF6, ORF7a, ORF7b and ORF9c, but significantly reduced in ORF8-expressing cells (Figure 2B). On their behalf, IL1b was significantly overexpressed in cells expressing ORF3d, ORF6, ORF8 and ORF9c (Figure 2C). Surprisingly, ORF8-expressing cells showed the highest increase in IL1b expression, but the lower in IL8 expression (Figure 2B and 2C), suggesting a compensatory mechanism between both interleukins signaling that merits further research. However, cells infected with SARS-CoV-2 virus showed a significant increase in IL11 and IL8 expression, but not in IL6 or IL1b. According to these results, other studies have also described an IL11 increase in A549-ACE2 and Calu-3 lung epithelial cells infected with SARS-CoV-2 ^10,58^ and COVID-19 lung biopsies ^10,59^. In addition, both cells and lung biopsies showed a similar profibrotic response to SARS-CoV-2 infection, as A549 cells expressing ORF6, ORF8, ORF9b and ORF9c ^10^. By contrast, it has been reported an increase in IL6 expression in other lung epithelial cells, such as Calu3-2B4^15^, or lungs of infected mice^12^. Our results agree with the epithelial model used in this study, and may indicate a possible interaction between IL11 and IL8 interleukins resulting in IL1b blockade. Indeed, it has been reported that IL1b increased levels during infection is induced by inflammasome activation, not by IL1b mRNA overexpression ^60–62^. Moreover, previous results from our group showed an increase in IL1b expression in ORF9b-expressing cells treated with an IL11 receptor inhibitor (Bazedoxifene), suggesting a possible crosslink between IL11 and IL1b signaling pathways ^10^.

Cytokine levels detected by ELISA showed very low levels in IL6 release by ORF-A549 cells, and high levels of IL11 and IL8 levels in most of transduced cells (Figure 3A, 3B and 3C). These data are consistent with those observed by RT-qPCR (Figure 2A, 2B and 2C). Since it has been described that human alveolar and bronchial epithelial-like cells produce IL11 ^54^, it makes sense that our A549 transduced cells show higher levels of IL11 release (Figure 3A). That might also happen in the lungs of the patients. Indeed, high levels of IL11RA (interleukin 11 receptor subunit alpha) mRNA and protein, but not IL11, have been detected in serum from COVID-19 patients ^63^. In fact, those authors suggest to use IL11RA expression potential discriminative ability among healthy, mild and severe SARS-CoV-2 cases ^63^.

The results observed in this study correlate with the inflammatory context induced by SARS-CoV-2 infection, with an excessive release of proinflammatory cytokines. The use of several anti-inflammatory drugs, such as Dexamethasone (a strong corticoid), to counteract the cytokine storm occurring during mild and severe disease, and the suggested use of IL11 as a therapeutic target in several diseases ^31,32^, led us to treat our A549 transduced cells with potential anti-inflammatory compounds. Since we found IL11 was completely inhibited by Dexamethasone, we decided to assay the suitability of this experimental system to evaluate compounds with potential anti-inflammatory activity towards SARS-CoV-2-induced inflammatory responses. For that purpose, tegument- and somatic-enriched protein extracts of newly excysted juveniles of *Fasciola hepatica* (FhNEJ-Teg and FhNEJ-Som, respectively) were tested. It is known that *F. hepatica*, and generally all parasitic helminths, have evolved sophisticated mechanisms that allow them to escape from host immunity ^36,37^. Such mechanisms induce an overall state of immune down-regulation that has been suggested to mitigate inflammatory processes, including those triggered by SARS-CoV-2 infection ^38–43^. Based on this knowledge, we treated ORF10-transduced cells with FhNEJ-Teg or FhNEJ-Som at different concentrations and measured IL-11 release and cell viability 24 hours after treatment (Figure 4). Due to technical challenges encountered during protein extraction of the somatic fraction of FhNEJ, the maximum concentration of FhNEJ-Som that could be added to cells without causing substantial toxicity was generally lower than that used for FhNEJ-Teg. This said, and despite the significant decrease in cell viability due to toxicity of FhNEJ-Som buffer compounds, results showed that this antigenic extract contains proteins that inhibit IL11 secretion by ORF10-transduced cells. These results align with the well-established ability of helminth parasites, including *F. hepatica*, to suppress inflammatory responses in theirmammalian hosts ^36,37,64^ and nicely complement previous findings by us, which demonstrated that FhNEJ-Teg contains proteins with antiviral potential that inhibit infection efficiency of both SARS-CoV-2 pseudotyped and live viral particles ^46^. Most importantly, our data demonstrate that the cellular platform developed in this study (INCEPLAT) was a valuable tool for identifying novel compounds with anti-inflammatory potential.

## Supporting information

List of primers

## AUTHOR CONTRIBUTIONS

Conceptualization: MM and BDLA

Methodology: all authors

Investigation: all authors

Funding acquisition: MM, MSL and JJG

Writing – original draft: BDLA, JS, MSL and MM

Writing – review & editing: BDLA, MM and SZ

## REFERENCES

1. Steiner S, Kratzel A, Barut GT, et al. SARS-CoV-2 biology and host interactions. Nat Rev Microbiol. 2024;22(4):206–225. doi:10.1038/s41579-023-01003-z

2. Gordon DE, Jang GM, Bouhaddou M, et al. A SARS-CoV-2 protein interaction map reveals targets for drug repurposing. Nature. 2020;583(7816):459-468. doi:10.1038/s41586-020-2286-9

3. Jungreis I, Nelson CW, Ardern Z, et al. Conflicting and ambiguous names of overlapping ORFs in the SARS-CoV-2 genome: A homology-based resolution. Virology. 2021;558:145–151. doi:10.1016/j.virol.2021.02.013

4. Kim D, Lee JY, Yang JS, Kim JW, Kim VN, Chang H. The Architecture of SARS-CoV-2 Transcriptome. Cell. 2020;181(4):914–921.e10. doi:10.1016/j.cell.2020.04.011

5. Li X, Hou P, Ma W, et al. SARS-CoV-2 ORF10 suppresses the antiviral innate immune response by degrading MAVS through mitophagy. Cell Mol Immunol. 2022;19(1):67–78. doi:10.1038/s41423-021-00807-4

6. Ramachandran K, Maity S, Muthukumar AR, et al. SARS-CoV-2 infection enhances mitochondrial PTP complex activity to perturb cardiac energetics. iScience. 2022;25(1):103722. doi:10.1016/j.isci.2021.103722

7. Xia H, Cao Z, Xie X, et al. Evasion of Type I Interferon by SARS-CoV-2. Cell Rep. 2020;33.

8. Zhang Y, Chen Y, Li Y, et al. The ORF8 protein of SARS-CoV-2 mediates immune evasion through down-regulating MHC-Ι. PNAS. 2021;118.

9. García-García T, Fernández-Rodríguez R, Redondo N, et al. Impairment of antiviral immune response and disruption of cellular functions by SARS-CoV-2 ORF7a and ORF7b. iScience. 2022;25(11). doi:10.1016/j.isci.2022.105444

10. López-Ayllón BD, de Lucas-Rius A, Mendoza-García L, et al. SARS-CoV-2 accessory proteins involvement in inflammatory and profibrotic processes through IL11 signaling. Front Immunol. 2023;14. doi:10.3389/fimmu.2023.1220306

11. López-Ayllón BD, Marin S, Fariñas Fernández M, et al. Metabolic and mitochondria alterations induced by SARS-CoV-2 accessory proteins ORF3a, ORF9b, ORF9c and ORF10. J Med Virol. Published online 2024.

12. Bello-Perez M, Hurtado-Tamayo J, Mykytyn AZ, et al. SARS-CoV-2 ORF8 accessory protein is a virulence factor. mBio. 2023;14(5). doi:10.1128/mbio.00451-23

13. Jiang H wei, Zhang H nan, Meng Q feng, et al. SARS-CoV-2 Orf9b suppresses type I interferon responses by targeting TOM70. Cell Mol Immunol. 2020;17(9):998–1000. doi:10.1038/s41423-020-0514-8

14. Mcgrath ME, Xue Y, Dillen C, et al. SARS-CoV-2 variant spike and accessory gene mutations alter pathogenesis. Published online 2022. doi:10.1073/pnas

15. Wang L, Guzman M, Muñoz-Santos D, et al. Cell type dependent stability and virulence of a recombinant SARS-CoV-2, and engineering of a propagation deficient RNA replicon to analyze virus RNA synthesis. Front Cell Infect Microbiol. 2023;13. doi:10.3389/fcimb.2023.1268227

16. Silvas JA, Vasquez DM, Park JG, et al. Contribution of SARS-CoV-2 Accessory Proteins to Viral Pathogenicity in K18 Human ACE2 Transgenic Mice. J Virol. 2021;95(17). doi:10.1128/jvi.00402-21

17. Miorin L, Kehrer T, Sanchez-Aparicio MT, et al. SARS-CoV-2 Orf6 hijacks Nup98 to block STAT nuclear import and antagonize interferon signaling. Proceedings of the National Academy of Sciences. 2020;117(45):28344–28354. doi:10.1073/pnas.2016650117

18. Coomes EA, Haghbayan H. Interleukin-6 in Covid-19: A systematic review and meta-analysis. Rev Med Virol. 2020;30(6):1–9. doi:10.1002/rmv.2141

19. Rabaan AA, Al-Ahmed SH, Muhammad J, et al. Role of inflammatory cytokines in covid-19 patients: A review on molecular mechanisms, immune functions, immunopathology and immunomodulatory drugs to counter cytokine storm. Vaccines (Basel). 2021;9(5). doi:10.3390/vaccines9050436

20. Montazersaheb S, Hosseiniyan Khatibi SM, Hejazi MS, et al. COVID-19 infection: an overview on cytokine storm and related interventions. Virol J. 2022;19(1):92. doi:10.1186/s12985-022-01814-1

21. Cook SA, Schafer S. Hiding in Plain Sight: Interleukin-11 Emerges as a Master Regulator of Fibrosis, Tissue Integrity, and Stromal Inflammation. Annu Rev Med. 2020;71(1):263–276. doi:10.1146/annurev-med-041818-011649

22. Metcalfe RD, Putoczki TL, Griffin MDW. Structural Understanding of Interleukin 6 Family Cytokine Signaling and Targeted Therapies: Focus on Interleukin 11. Front Immunol. 2020;11. doi:10.3389/fimmu.2020.01424

23. Giraldez MD, Carneros D, Garbers C, Rose-John S, Bustos M. New insights into IL-6 family cytokines in metabolism, hepatology and gastroenterology. Nat Rev Gastroenterol Hepatol. 2021;18(11):787–803. doi:10.1038/s41575-021-00473-x

24. Rose-John S. Interleukin-6 Family Cytokines. Cold Spring Harb Perspect Biol. 2018;10(2):a028415. doi:10.1101/cshperspect.a028415

25. Widjaja AA, Chothani SP, Cook SA. Different roles of interleukin 6 and interleukin 11 in the liver: implications for therapy. Hum Vaccin Immunother. 2020;16(10):2357–2362. doi:10.1080/21645515.2020.1761203

26. Elias JA, Zheng T, Einarsson O, et al. Epithelial interleukin-11: Regulation by cytokines, respiratory syncytial virus, and retinoic acid. Journal of Biological Chemistry. 1994;269(35):22261–22268. doi:10.1016/s0021-9258(17)31785-4

27. Corne JM, Holgate ST. Mechanisms of virus induced exacerbations of asthma. Thorax. 1997;52(4):380–389. doi:10.1136/thx.52.4.380

28. Einarsson O, Geba GP, Zhu Z, Landry M, Elias J. Interleukin-11: Stimulation In Vivo and In Vitro by Respiratory Viruses and Induction of Airways Hyperresponsiveness. J Clin Invest. 1996;97:915–924.

29. Widjaja AA, Viswanathan S, Jinrui D, et al. Molecular Dissection of Pro-Fibrotic IL11 Signaling in Cardiac and Pulmonary Fibroblasts. Front Mol Biosci. 2021;8. doi:10.3389/fmolb.2021.740650

30. Ng B, Dong J, D’Agostino G, et al. Interleukin-11 is a therapeutic target in idiopathic pulmonary fibrosis. Sci Transl Med. 2019;11(511). doi:10.1126/scitranslmed.aaw1237

31. Ng B, Xie C, Su L, et al. IL11 (Interleukin-11) Causes Emphysematous Lung Disease in a Mouse Model of Marfan Syndrome. Arterioscler Thromb Vasc Biol. 2023;43(5).

32. Cook SA. Understanding interleukin 11 as a disease gene and therapeutic target. Biochemical Journal. 2023;480(23):1987–2008. doi:10.1042/BCJ20220160

33. Sinha S, Rosin NL, Arora R, et al. Dexamethasone modulates immature neutrophils and interferon programming in severe COVID-19. Nat Med. 2022;28(1):201–211. doi:10.1038/s41591-021-01576-3

34. Tomazini BM, Maia IS, Cavalcanti AB, et al. Effect of Dexamethasone on Days Alive and Ventilator-Free in Patients With Moderate or Severe Acute Respiratory Distress Syndrome and COVID-19. JAMA. 2020;324(13):1307. doi:10.1001/jama.2020.17021

35. Siles-Lucas M, Becerro-Recio D, Serrat J, González-Miguel J. Fascioliasis and fasciolopsiasis: Current knowledge and future trends. Res Vet Sci. 2021;134:27–35. doi:10.1016/j.rvsc.2020.10.011

36. Maizels RM. Regulation of immunity and allergy by helminth parasites. Allergy: European Journal of Allergy and Clinical Immunology. 2020;75(3):524–534. doi:10.1111/all.13944

37. Maizels RM, McSorley HJ. Regulation of the host immune system by helminth parasites. Journal of Allergy and Clinical Immunology. 2016;138(3):666–675. doi:10.1016/j.jaci.2016.07.007

38. Ademe M, Girma F. The influence of helminth immune regulation on covid-19 clinical outcomes: Is it beneficial or detrimental? Infect Drug Resist. 2021;14:4421–4426. doi:10.2147/IDR.S335447

39. Bradbury RS, Piedrafita D, Greenhill A, Mahanty S. Will helminth co-infection modulate COVID-19 severity in endemic regions? Nat Rev Immunol. 2020;20(6):342. doi:10.1038/s41577-020-0330-5

40. Siles-Lucas M, González-Miguel J, Geller R, Sanjuan R, Pérez-Arévalo J, Martínez-Moreno A. Potential Influence of Helminth Molecules on COVID-19 Pathology. Trends Parasitol. Published online 2020.

41. Whitehead B, Christiansen S, Østergaard L, Nejsum P. Helminths and COVID-19 susceptibility, disease progression, and vaccination efficacy. Trends Parasitol. 2022;38(4):277–279. doi:10.1016/j.pt.2022.01.007

42. Wolday D, Gebrecherkos T, Arefaine ZG, et al. Effect of co-infection with intestinal parasites on COVID-19 severity: A prospective observational cohort study. EClinicalMedicine. 2021;39. doi:10.1016/j.eclinm.2021.101054

43. Fonte L, Acosta A, Sarmiento ME, Ginori M, García G, Norazmi MN. COVID-19 Lethality in Sub-Saharan Africa and Helminth Immune Modulation. Front Immunol. 2020;11. doi:10.3389/fimmu.2020.574910

44. Cruz JLG, Becares M, Sola I, Oliveros JC, Enjuanes L, Zúñiga S. Alphacoronavirus Protein 7 Modulates Host Innate Immune Response. J Virol. 2013;87(17):9754–9767. doi:10.1128/JVI.01032-13

45. Serrat J, Becerro-Recio D, Torres-Valle M, et al. Fasciola hepatica juveniles interact with the host fibrinolytic system as a potential early-stage invasion mechanism. PLoS Negl Trop Dis. 2023;17(4). doi:10.1371/journal.pntd.0010936

46. Serrat J, Francés-Gómez C, Becerro-Recio D, González-Miguel J, Geller R, Siles-Lucas M. Antigens from the Helminth Fasciola hepatica Exert Antiviral Effects against SARS-CoV-2 In Vitro. Int J Mol Sci. 2023;24(14). doi:10.3390/ijms241411597

47. Cwiklinski K, Robinson MW, Donnelly S, Dalton JP. Complementary transcriptomic and proteomic analyses reveal the cellular and molecular processes that drive growth and development of Fasciola hepatica in the host liver. BMC Genomics. 2021;22(1). doi:10.1186/s12864-020-07326-y

48. López-Ayllón BD, de Lucas-Rius A, Mendoza-García L, et al. SARS-CoV-2 accessory proteins involvement in inflammatory and profibrotic processes through IL11 signaling. Front Immunol. 2023;14. doi:10.3389/fimmu.2023.1220306

49. Grant E Keeney et al. Dexamethasone for acute asthma exacerbations in children: a meta-analysis. Pediatrics. 2014;Mar;133(3):493–499.

50. P M Irving RBGMPSPRG. Review article: appropriate use of corticosteroids in Crohn’s disease. Aliment Pharmacol Ther. 2008;15;27(6):528–529.

51. Basil A. Stoll F.F.R. Dexamethasone in advanced breast cancer. *Basil A Stoll FFR*, MRCS. Published online 1960:1074–1080.

52. Joseph L. Hollander MD. Clinical use of Dexamethasone role in treatment of patients with asrthritis. JAMA. 1960;172(4):306–310.

53. Ng B, Dong J, D’Agostino G, et al. Interleukin-11 is a therapeutic target in idiopathic pulmonary fibrosis. Sci Transl Med. 2019;11(511). doi:10.1126/scitranslmed.aaw1237

54. Elias JA, Zheng T, Einarsson O, et al. Epithelial interleukin-11: Regulation by cytokines, respiratory syncytial virus, and retinoic acid. Journal of Biological Chemistry. 1994;269(35):22261–22268. doi:10.1016/s0021-9258(17)31785-4

55. Sánchez-Cabo F, Fuster V, Silla-Castro JC, et al. Subclinical atherosclerosis and accelerated epigenetic age mediated by inflammation: a multi-omics study. Eur Heart J. 2023;44(29):2698–2709. doi:10.1093/eurheartj/ehad361

56. Unamuno X, Gómez-Ambrosi J, Ramírez B, et al. NLRP3 inflammasome blockade reduces adipose tissue inflammation and extracellular matrix remodeling. Cell Mol Immunol. 2021;18(4):1045–1057. doi:10.1038/s41423-019-0296-z

57. Kraan MC, Patel DD, Haringman JJ, et al. The Development of Clinical Signs of Rheumatoid Synovial Inflammation Is Associated with Increased Synthesis of the Chemokine CXCL8 (Interleukin-8). http://arthritis-research.com/content/3/1/065

58. Blanco-Melo D, Nilsson-Payant BE, Liu WC, et al. Imbalanced Host Response to SARS-CoV-2 Drives Development of COVID-19. Cell. 2020;181(5):1036–1045.e9. doi:10.1016/j.cell.2020.04.026

59. Wang S, Yao X, Ma S, et al. A single-cell transcriptomic landscape of the lungs of patients with COVID-19. Nat Cell Biol. 2021;23(12):1314–1328. doi:10.1038/s41556-021-00796-6

60. Diarimalala RO, Wei Y, Hu D, Hu K. Inflammasomes during SARS-CoV-2 infection and development of their corresponding inhibitors. Front Cell Infect Microbiol. 2023;13. doi:10.3389/fcimb.2023.1218039

61. Freeman TL, Swartz TH. Targeting the NLRP3 Inflammasome in Severe COVID-19. Front Immunol. 2020;11. doi:10.3389/fimmu.2020.01518

62. Rodrigues TS, Zamboni DS. Inflammasome activation by SARS-CoV-2 and its participation in COVID-19 exacerbation. Curr Opin Immunol. 2023;84. doi:10.1016/j.coi.2023.102387

63. Agwa SHA, Elghazaly H, Meteini MS El, et al. In silico identification and clinical validation of a novel long non-coding rna/mrna/mirna molecular network for potential biomarkers for discriminating sars cov-2 infection severity. Cells. 2021;10(11). doi:10.3390/cells10113098

64. Ryan S, Shiels J, Taggart CC, Dalton JP, Weldon S. Fasciola hepatica-Derived Molecules as Regulators of the Host Immune Response. Front Immunol. 2020;11. doi:10.3389/fimmu.2020.02182

