## Supplementary material for "Inflammation cellular platform (INCEPLAT) for testing anti-inflammatory compounds for SARS-CoV-2": Supp Fig 1S

**Figure S1**
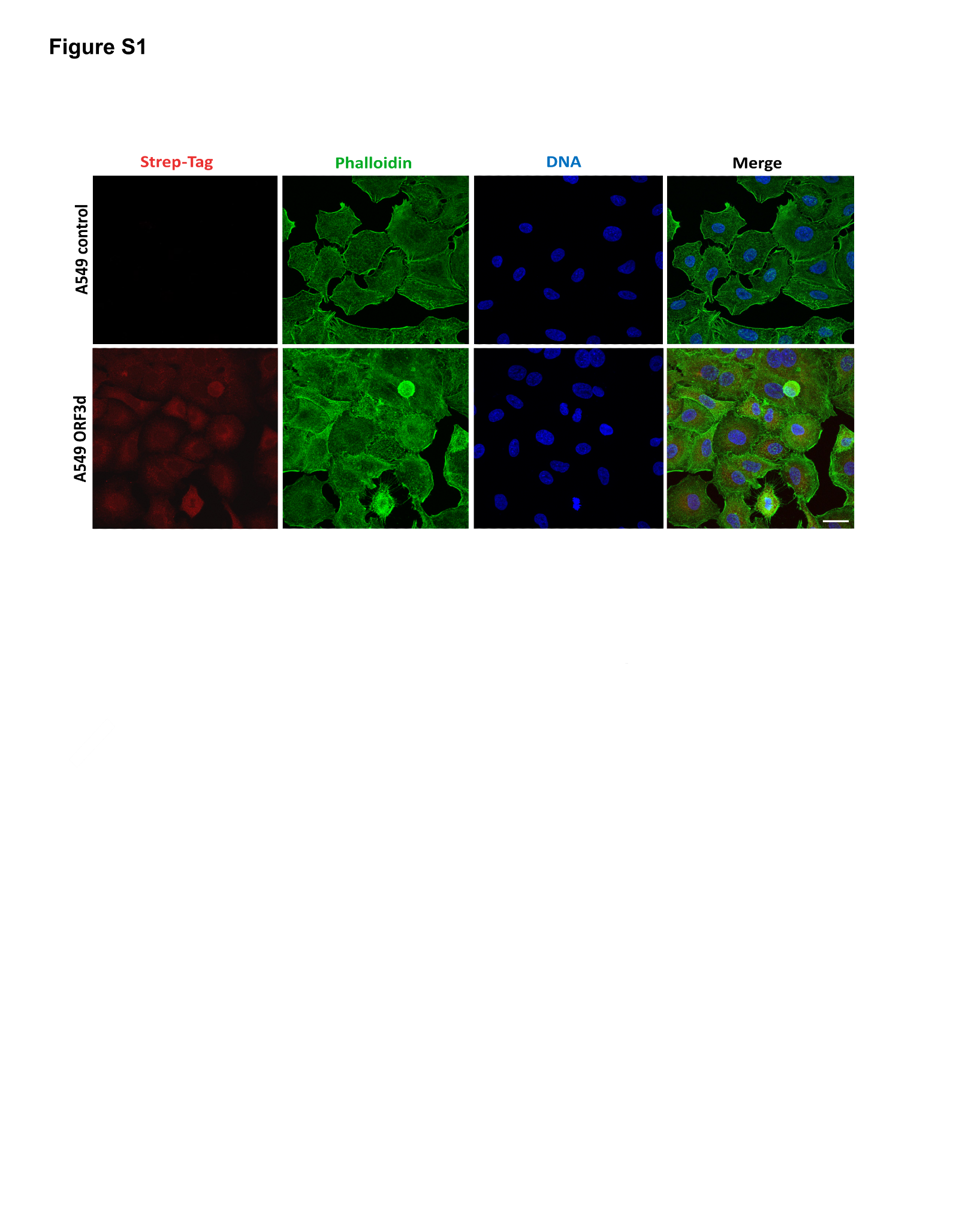

**Figure S1**. Representative confocal images of A549 cells expressing SARS-CoV-2 ORF3d accessory protein (objective 63x, scale bar 25 µm).
