## Supplementary material for "Inflammation cellular platform (INCEPLAT) for testing anti-inflammatory compounds for SARS-CoV-2": List of primers

**Table S1. List of primers used for RT-qPCR.**

| **Gene** | **Forward primer (5'-3')** | **Reverse primer (5'-3')** | **Reference** |
| --- | --- | --- | --- |
| IL11 | GAAACAGCAGGCTACAAAACCACT | ACCTCTCTCCTTTGACCTGGAGAC | López-Ayllón BD, et al 2023 |
| IL8 | GAAGAAACCACCGGAAGGAA | CAAAACTGCACCTTCACACA | This work |
| IL1β | ACAGATGAAGTGCTCCTTCC | CGGCCTGCCTGAAGCCCTTG | López-Ayllón BD, et al 2023 |
| IL6 | GATGGATGCTTCCAATCTGGAT | CAGCTCTGGCTTGTTCCTCACT | This work |
| GAPDH | TGGGTGTGAACCATGAGAAG | TGGCAGTGATGGCATGGAC | López-Ayllón BD, et al 2023 |
